# Temperature-Induced Uncoupling of Cell Cycle Regulators

**DOI:** 10.1101/2020.02.11.943266

**Authors:** Hanieh Falahati, Woonyung Hur, Stefano Di Talia, Eric F. Wieschaus

**Author notes:** For correspondence (EFW). HHMI Life Sciences Associate, Departments of Neuroscience and of Cell Biology, Kavli Institute for Neuroscience, Yale University School of Medicine.

## Abstract

While feedback loops are essential for robustness in signaling systems, they make discerning the role of individual components challenging. Here we introduce temperature as a powerful perturbation method for uncoupling enzymatic processes, by exposing the differential sensitivity of limiting reactions to temperature due to their activation energies. Using this method, we study the sensitivity to temperature of different cell cycle events of early fly embryos. While the subdivision of cell cycle steps is conserved across a wide range of temperatures (5-35°C), the transition into prometaphase exhibits the most sensitivity, arguing that it has a different mechanism of regulation. Using a biosensor, we quantify the activity of Cdk1 and show that the activation of Cdk1 drives entry into prometaphase but is not required for earlier events. In fact, chromosome condensation can still occur when Cdk1 is inhibited pharmacologically. These results demonstrate that different kinases are rate-limiting for different steps of mitosis.

## Introduction

Understanding how sequences of cellular transitions are regulated remains a fundamental, yet poorly understood, biological question. During cell cycle progression, for example, it remains unclear whether the various morphological transitions between interphase and mitosis reflect the accumulation pattern of a single rate limiting regulator, or whether each step is associated with its own distinct regulatory components. The complex feedbacks that characterize genetic networks (***Morgan, 2007***; ***Lindqvist et al., 2009***; ***Heim et al., 2016***) make it difficult to define roles of single components using null mutations and has led to conflicting conclusions on the relative importance of the various kinases involved. Here, we use subtle perturbations in temperature as an alternative to distinguish the mechanisms regulating different transitions. Our approach is centered on the idea that the behavior of different enzymes at varying temperatures is characterized by different scaling relationships. Thus, measuring the dependence of the timing of cellular events on temperature could reveal their underlying regulatory dynamics.

## Results and Discussion

We used temperature to study the regulation of cell cycle in early *D. melanogaster* embryos. The cell cycle proceeds through a strict sequence of events - DNA replication and centrosome doubling are followed by chromatin condensation, movement of the chromosomes to a single metaphase plate and their eventual separation at mitosis that can be easily scored in living embryos. The timely order with which these events occur is pivotal for the successful transfer of genetic material to daughter cells and several regulatory mechanisms have been linked to an autonomous oscillator centered on the activity of Cdk1 (***Pomerening et al., 2003***; ***Morgan, 2007***; ***Ferree and Di Talia, 2018***). Whether individual mitotic events are triggered at specific levels of activity of Cdk1 or whether other mitotic kinases, like Polo and Aurora B, coupled to Cdk1 play more direct roles in regulating mitotic processes remains to be elucidated. We reasoned that precise measurements of the temperature dependency of mitotic processes might reveal novel insights. For a given enzyme, in first approximation, temperature (*T*) affects its rate (*k*) by Arrhenius equation (***Arrhenius, 1915***):

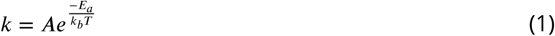

where *A, E*_*a*_ and *k*_*b*_ are the pre-exponential factor, activation energy, and Boltzmann constant, respectively. Thus, if Cdk1 was rate-limiting for all mitotic processes, we would expect that these processes will show a similar dependency on temperature. In contrast, if different mitotic kinases contributed to the regulation of the timing of mitotic processes, it is likely that we would observe different Arrhenius dependencies. We propose to exploit this differential sensitivity to uncouple cell cycle processes and evaluate the contributions of Cdk1 in regulation of the minimal cell cycles of early *Drosophila* embryos.

The first 13 divisions in *Drosophila* embryos occur synchronously through rapid rounds of DNA synthesis (S-phase) and nuclear mitosis (***Rabinowitz, 1941***; ***Farrell et al., 2012***). For the purpose of this study, we mostly focus on nuclear cycle 11 (NC11) due to the ease of imaging. To examine the effect of temperature on cell cycle steps, we used a microfluidic device to control the temperature of the embryos (between 5-35°C) while performing live imaging (***Lucchetta et al., 2005***; ***Falahati and Wieschaus, 2017***). While lowering the temperature to a minimum of 5°C increases the duration of NC11 (from 10m:30s±15s at 22°C to 92min±12min at 6°C, ***Figure 1****A*), it does not change the examined events characteristic of each cell cycle step. In particular, as depicted in ***Figure 1****B* and ***Video 1*** for embryos developing at 22 and 7°C, there are no detectable differences in the large-scale chromatin structure and chromosomal movements, as visualized by fluorescently-tagged Histone H2A variant (H2Av). Interestingly, we also found no detectable irregularities in centrosome cycle at low temperatures. To visualized the microtubule organization, we imaged a microtubule associated protein (MAP), Jupiter-GFP (***Karpova et al., 2006***), and quantified the distances of the centriole pairs (***Figure 1****B*, ***Figure 1***–***Figure Supplement 1***, and ***Video 2***). Similar to systems with cold stabilizing MAPs (***Denarier et al., 1998***), no reduction is detectable in the density of microtubules in fly embryos at 8°C. Finally, between 22 and 5°C, a conserved pattern is observed for the changes in the nuclear envelop integrity, and also for the distribution of the DNA replication machinery at different steps of cell cycle, as visualized, respectively, by fluorescently-tagged nuclear protein Fibrillarin (***Falahati et al., 2016***), and replication machinery scaffold protein PCNA (***Farrell et al., 2012***; ***Blythe and Wieschaus, 2016***) (***Figure 1****B*, ***Figure 1***–***Figure Supplement 2***). However, this conservation of cell cycle events does not hold at 35°C, as defects in chromosome segregation can be noticed at this temperature (***Figure 1***–***Figure Supplement 4***), which is consistent with previous reports of induction of heat-shock mechanisms at this temperature (***Dura, 1981***).

**Figure 1.**
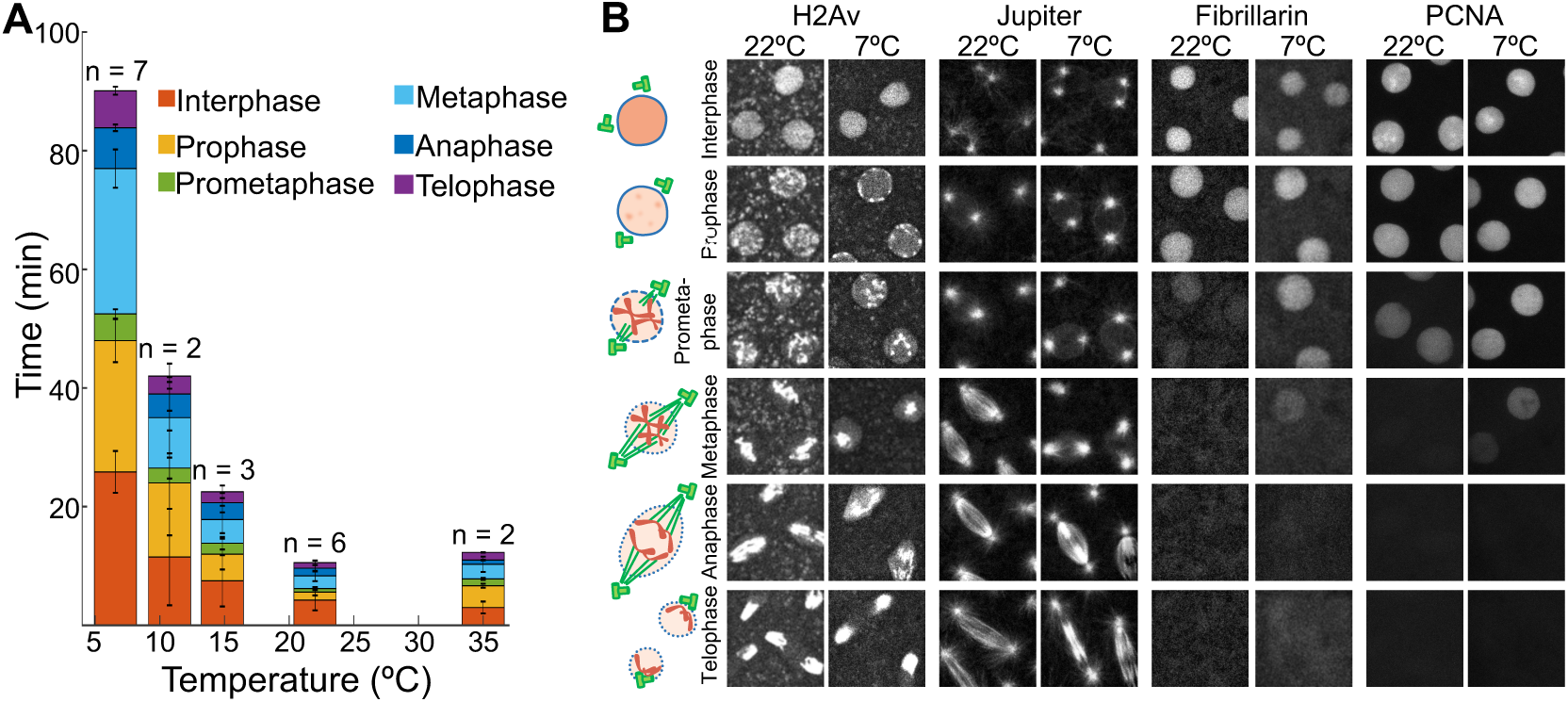
Effect of temperature on cell cycle events. **A** The total length of different steps of cell cycle were calculated based on the subcellular dynamics of H2Av. **B** The following fluorescently-tagged proteins were used as a proxy for different cell cycle events: H2Av for cell cycle steps; Jupiter for centrosome cycle; the nuclear protein, Fibrillarin, for nuclear envelope breakdown and reformation; and PCNA for DNA replication. Images are 25× 25*μm*. **Figure 1-Figure supplement 1**. A schematic of the micro2uidic device used for controling the temperature. **Figure 1-Figure supplement 2**. Distance of centrosomes at 22 and 8°*C*. **Figure 1-Figure supplement 3**. Quanti1cation of nuclear import and DNA replication at 22 and 8°*C*. **Figure 1-Figure supplement 4**. Development of NC11 embryos at 35°*C*.

**Video 1. The scaling of the length of cell cycle steps with temperature.** Two NC11 embryos developing at 22°C (Top) and 7°C (Bottom) are shown. The movies start at the beginning of interphase for both embryos. The frame-rates are set such that the initiation and end of interphase would appear at the same time in the movies of both embryos. Single z plane of 82.04× 40.98*μm* is shown for each.

**Video 2. Spindle formation at 7**°**C.** An NC11 embryo co-expressing H2Av-RFP (magenta) and GFP-tagged microtubule binding protein, Jupiter-GFP, (green) is shown at 7°C. Each time point is a maximum projection of 82.04× 40.98*μm*.

In addition to the qualitative similarities, we examined whether the changes in the length of NC steps follows the Arrhenius equation between 22-5°C. We calculated the rate at which each NC step progresses as the reciprocal of its duration, and plotted the log of the measured rates as a function of temperature 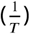. As depicted in ***Figure 2****A-B*, the rates of all NC steps between 5-22°C fit well with the Arrhenius equation (*r*^2^ ⩾ 0.82). A similar Arrhenius-dependency was also reported for the rate of completion of the first embryonic cycle of *C. elegans* at this temperature range (***Begasse et al., 2015***), and also for the total developmental time of *Drosophila* species between 17.5 and 27.5°C (***Kuntz and Eisen, 2014***). Since for each NC step, the dependency is linear between 5-22°C, the slope can be used to calculate the activation energies associated with each process (***Figure 2****C*). The *E*_*a*_ values obtained are within the range of other reported biological reactions (***Raven and Geider, 1988***; ***Gillooly et al., 2001***; ***Dell et al., 2011***).

**Figure 2.**
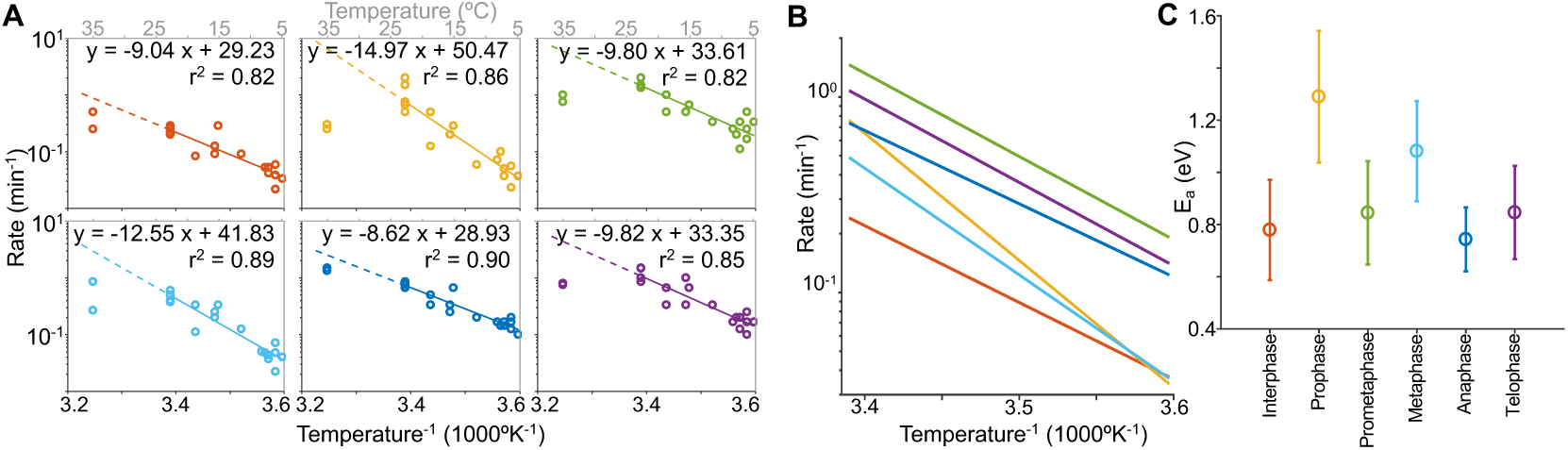
Arrhenius-dependence of cell cycle steps. **A** and **B** The Arrhenius plot for each of the cell cycle steps is depicted, with a line fitted to the linear region. A comparison of the fitted lines is shown in **B. C** shows the calculated *E*_*a*_ for each cell cycle step. Error bars: 95% confidence intervals.

Interestingly, within the linear regime, the Arrhenius plot of prophase temperature-response has the steepest slope, indicating that reductions in temperature have a much more significant impact on prophase duration than on the length of other steps of the cell cycle. This, in turn, suggests that initiation of prometaphase may be determined by a regulatory enzyme with an *E*_*a*_ higher than the regulators of other cell cycle steps. The prolonged prophase at low temperatures is biologically significant, resulting in hypercondensed chromosomes that decorate the nuclear envelope, while awaiting the delayed chromosome congression (***Figure 1****B*, ***Video 1***) and a delayed nuclear envelope breakdown, as visualized by redistribution of the nuclear protein, RFP-Fibrillarin (***Figure 1****B*, ***Figure 1***–***Figure Supplement 2****A*).

These results are suggestive of two main points: 1. Entry into prometaphase is regulated by an enzyme that is the most sensitive to temperature, with the highest *E*_*a*_, compared to other cell cycle regulators; and 2. Earlier NC events are not regulated by this most temperature sensitive enzyme. To characterize the enzyme driving the entry into prometaphase, we employed a Förster resonance energy transfer (FRET)-based sensor to monitor the activity of Cdk1 in embryos developing at different temperatures (***Gavet and Pines, 2010***; ***Deneke et al., 2016***). ***Figure 3****A* and ***Figure 3***–***Figure Supplement 1*** show the Cdk1 activity of NC11 embryos developing at 22 and 7°C. Although Cdk1 is known as the main regulator of mitotic progression, its activation at 7°C is only detectable at late prophase, immediately prior to the onset of prometaphase. Although exit from prophase is coupled to increased Cdk1 activity, entry, that is chromosome condensation, occurs when Cdk1 activity is much lower than that observed for prophase entry at higher temperatures. The observed differing dependence of prophase entry and exit was confirmed by examining cell cycle progression when Cdk1 activity is inhibited by roscovitine. Injection of roscovitine blocks exit from prophase, but had no effect on entry. Chromosomes initiate condensation (***Figure 3****C*, ***Video 3***) and ultimately show the same hypercondensation observed during the prolonged prophase at low temperatures (***Figure 3****C*). Together, these results indicate that at NC11, the activation of Cdk1 is necessary for entry into prometaphase, while earlier cell cycle events such as chromosome condensation can occur independently of Cdk1 activation.

**Figure 3.**
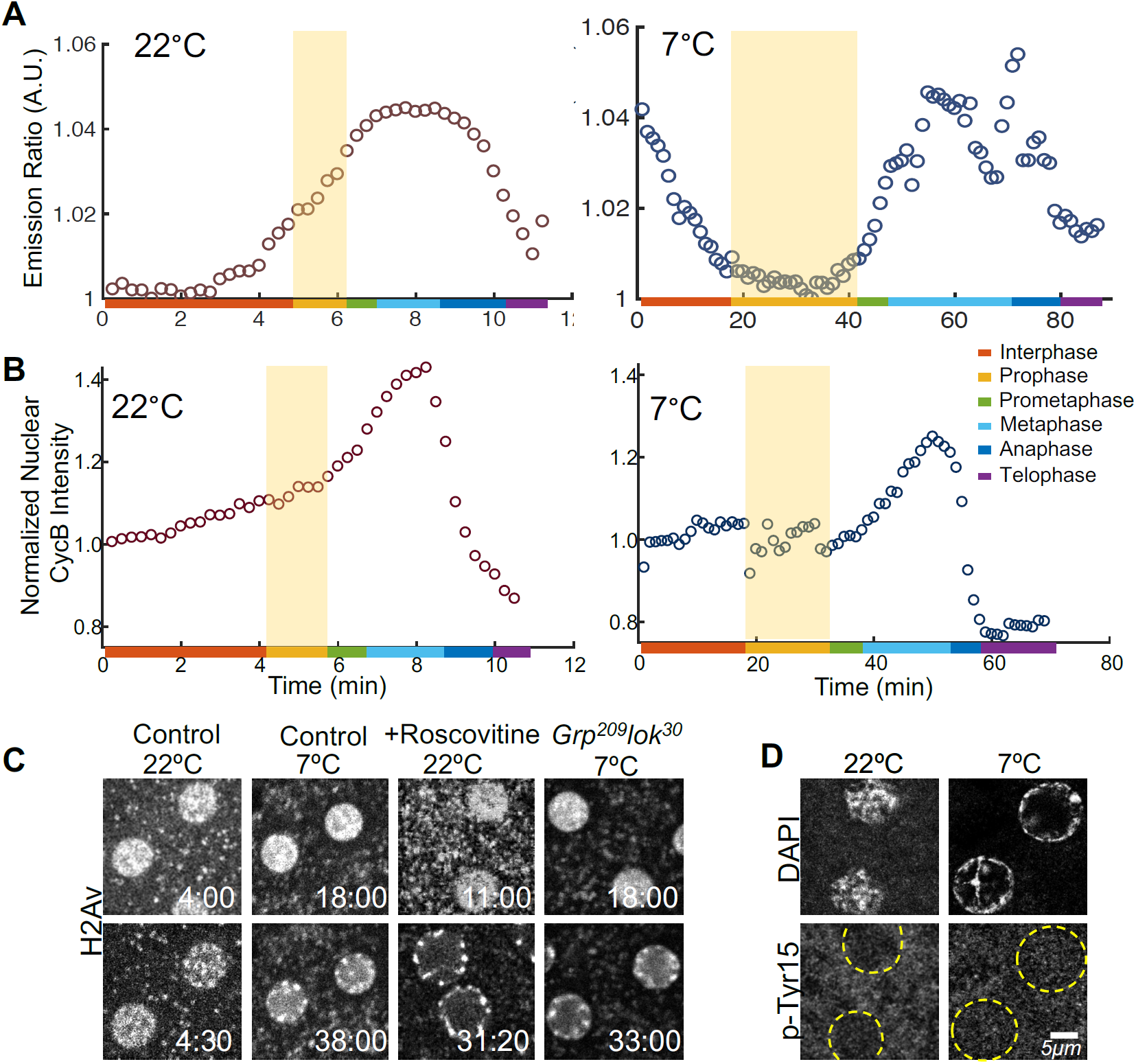
Cdk1 activity drives exit but not entry into prophase. **A** Cdk1 activity at 22 and 7°C is quantified using a FRET-based biosensor. **B** Normalized nuclear concentrations of CycB throughout NC11 at 22 and 7°C. Shaded areas show prophase. **C** Single plane images of WT untreated embryos (controls), WT embryos treated with roscovitine, and check-point mutant embryos (*Grp*^209^*lok*^30^), during early and late prophase at different temperatures. **D** Cdk1’s inhibitory p-Tyr15 is not detectable at any temperatures at NC11. Times in *min:sec*. **Figure 3-Figure supplement 1**. Other examples of Cdk1 emission ratio. **Figure 3-Figure supplement 2**. The relationship between Cdk1 activity and CycB accumulation show that CycB accumulation is rate limiting for Cdk1 activity.

**Video 3. Inhibition of Cdk1 prevents prometaphase but not prophase.** An NC11 embryo expressing H2Av-RFP injected with Cdk1 inhibitor, roscovitine is shown. The chromosome condensation happens in the absence of Cdk1 activity, but the embryo does not enter prometaphase. Each time-point is a single z plane of 58.12 × 36.30*μm*.

The altered cell cycle dynamics observed at 7°C raises one main question: What causes the high sensitivity of Cdk1 to temperature? It is unlikely that the delayed Cdk1 activation is caused by activation of DNA replication checkpoint, given that mutant embryos lacking the checkpoint components *grapes* and *loki* still exhibit chromosome hypercondensation at 7°C (***Figure 3****C*) and given previous reports that the inhibitory Tyr15 phosphorylation associated with checkpoint activation is not detectable in NC11 embryos at normal temperatures (***Edgar et al., 1994***). We confirm that the phosphorylation remains undetectable at 7°C (***Figure 3****D*). Instead the delayed rise in Ckd1 activity at 7°C appears to be related to a delayed increase in CycB levels. At 22°C, mean nuclear intensity of CycB measured using a GFP-CycB reporter increases monotonically from interphase, and sharply drops afterwards (***Figure 3****B*). At 7°C, the monotonic increase in the nuclear intensity of CycB is only detectable in prometaphase. At both temperatures, the kinetics of CycB accumulation parallels the rise in Cdk1 activity (***Figure 3***–***Figure Supplement 2***). The linear relationship between the nuclear accumulation of CycB and Cdk1 activity at both T’s suggests that in our experiments CycB accumulation is the temperature sensitive step that is rate limiting for the activation of Cdk1, a suggestion consistent with previous reports (***Edgar et al., 1994***; ***Pomerening et al., 2003***; ***Gavet and Pines, 2010***). The observed delay in Cdk1 activation at 7°C could either be through a slower synthesis rate of CycB or a delayed nuclear import. The sensitivity of our current reporters does not allow for distinguishing between these possibilities.

In conclusion, we have introduced temperature as a powerful perturbation tool to examine the limiting steps in complex biological processes. Using the combination of this method with optical biosensors and pharmacological approaches, we studied the regulatory mechanisms of cell cycle in fly embryos. Our results indicate that entry into prometaphase is regulated by Cdk1. However, earlier cell cycle events such as chromosome condensation and centrosome cycle can occur independently of Cdk1 activity, potentially by relying on other mitotic kinases such as Aurora and Polo (***Adams et al., 2001***; ***Giet and Glover, 2001***; ***McCleland and O’Farrell, 2008***). Consistent with this model, experiments in fly embryos using RNAi against mitotic cyclins show that the centriole cycle still continues in reduced Cdk1 activity (***Novak et al., 2016***). Together, these results suggest that the role of Cdk1 as the master regulator of cell cycle might not be as general as previously assumed.

## Methods and Materials

### Fabrication and utilization of the microfluidic device

The microfluidic device used here was fabricated as described previously (***Lucchetta et al., 2005***; ***Falahati and Wieschaus, 2017***). ***Figure 1***–***Figure Supplement 1*** shows the schematic of the device. It is comprised of two parts: The PDMS channel which is placed on top of a PDMS-coated cover glass. The embryos are mounted on the coated coverglass by placing a drop of heptane glue on top of the embryos. The excess heptane is swiftly wiped out. Approximately 20 embryos were mounted each time. After assembling the two parts, three pairs of small magnets were placed across the channels to lock the device in place and prevent leakage. A syringe pump was used to control the inward flow of water into the channels, and a second pump was placed at the outlet to apply suction and direct the flow through the channels. Since coordinating the speed of the two pumps is difficult, the suction pump was placed at full speed and an inlet for air was added to the channel close to the connection of the suction pump to compensate for the extra volume and prevent discontinuity in the flow of water passing through the embryos. The temperature of the fluid is measured by two thermometers placed at 0.5*cm* upstream the embryos. The temperatures were kept at ±1°C of the reported values.

### Staging of live embryos

The staging of live embryos was done using fluorescently-tagged H2Av. The beginning of interphase was denoted when the chromosomes decondense and histone is homogeneously distributed in the nucleus. The start of prophase is noted by reappearance of inhomogeneities in the distribution of histone, indicative of the beginning of chromosome condensation. Prometaphase starts with chromosomes’ migration toward the metaphase plate. The beginning of metaphase is when all chromosomes reach the metaphase plate. The separation of sister chromatids toward opposing poles determines the duration of anaphase, and chromosome decondensation occurs at telophase. The rate of each step shown in ***Figure 1*** is defined as the inverse of the time required for the completion of that particular step.

### Transgenic lines

Flies expressing Cdk1 FRET sensor (***Deneke et al., 2016***), PCNA-EGFP (***Blythe and Wieschaus, 2016***; ***Falahati and Wieschaus, 2017***), and RFP-Fibrillarin (***Falahati and Wieschaus, 2017***) were described previously. CycB-GFP fly stock is from Bloomington *Drosophila* Stock Center (BDSC 51568).

### Confocal imaging and image analysis

Embryos were imaged either using 63×HCX PL APO CS 1.4 NA oil-immersion objective on a Leica SP5 laser-scanning confocal microscope equipped with GaAsP ‘HyD’ detectors, or Leica SP8 confocal microscope, 20x/0.75 numerical aperture oil-immersion objective. All image analyses were performed with ImageJ (Rasband WS, ImageJ; National Institutes of Health; (1997-2008)) and MATLAB (MathWorks, Natick, MA) routines. To measure the changes in nuclear concentration of CycB, embryos coexpressing CycB-GFP and H2Av-RFP were imaged. The maximum projected images were used for quantification, and a nuclear mask was generated by applying a Gaussian filter and thresholding the H2Av-RFP signal. The mask was then used to measure the average nuclear intensity of CycB-GFP. This average intensity is then divided by its initial value in each cell cycle to normalize. To measure the distance between two centrosomes, embryos coexpressing H2Av-RFP and Jupiter-GFP were imaged at different temperatures. An ImageJ macro was developed to automatically segment the field of view to detect the area surrounding each nucleus. This segmentation was manually evaluated to fix the false positive/negatives. Then in each segmented area, the local maxima of the Jupiter-GFP signal were detected automatically and reconfirmed manually. Finally, the distance between the two centrosomes in each image was measured.

PCNA foci where quantified by measuring the local inhomogeneity of the GFP signal in the nucleus over time, as described previously (***Falahati and Wieschaus, 2017***). Maximum projected images were used and the homogeneity in the pixel intensities were measure by a customized GLCM method. To reduce the noise, the neighboring 5×5 pixels were averaged. The homogeneity was measured using MATLAB’s built-in function with the following equation:

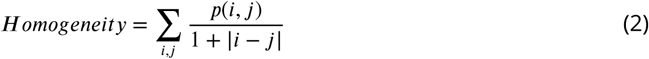

where *i* and *j* are the centers of intensity bins, and *p*(*i, j*) is the probability of two particular intensities being neighbors.

For FRET measurements, the fluorescent intensity of CFP and YFP were measured by averaging over the part of the embryo that is in the field of view at different conditions and different time-points, and the ratio of YFP/CFP is calculated.

### Fixation and immunostaining at different temperatures

The newly laid embryos were collected at 22°C for 90min and then incubated either at 22°C or at a cooling device set at 7°C for 30*min*. For embryos incubated at 7°C, the fixation process was carried out at 4°C, using pre-cooled fixative. The primary antibody used is rabbit Cdc2 phosphor-Tyr15 (IMG-668, IMGENEX).

### Injection of inhibitors

All the embryos were collected without using the halocarbon oil (this is to make sure that embryos are not protected from desiccation because of the oil), dechorionated in 50% bleach for 1 minute, desiccated in a chamber filled with Drierite (desiccant) for 8 minutes, and the drug was injected into the middle of the embryo using the Eppendorf Femtojet microinjector.

## Acknowledgments

We thank the Bloomington *Drosophila* Stock Center for fly stocks. We thank all members of E.F.W., S.D.T. and Schupbach laboratories the lively discussions. This work was in part supported by NIH (R01-GM 122936 to S.D.T). E.F.W. is an HHMI investigator. H.F. is an HHMI Life Sciences Associate.

---

**Figure 1–Figure supplement 1.**
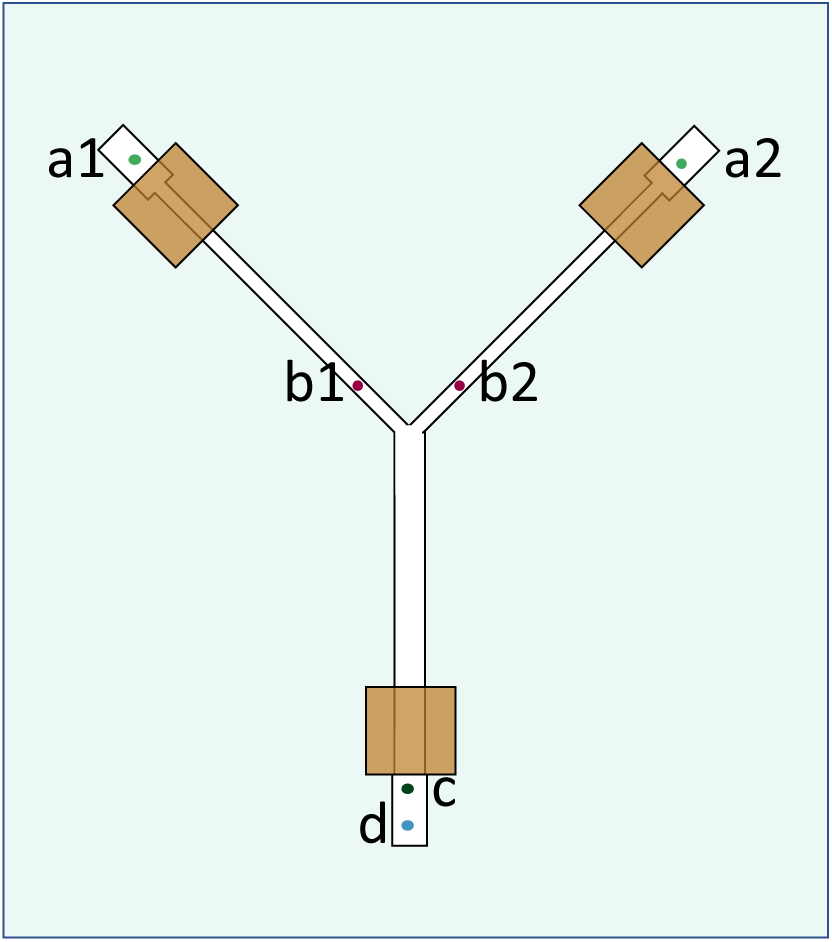
A schematic of the microfluidic device used for controling the temperature. The channels are shown in white. a1 and a2 are inlets that are connected to syringe pump that pushes the fluid into the channels. The two channels can be used for two different temperatures. b1 and b2 are connected to the thermometer for readout of the flow temperature. The suction pump is connected to c, and d is left open, to let the air flow inside the channel for compensation of the inward and outward flow to assure the continuity of the fluid passing over the embryos. Three pairs of magnets are used (squares in hazelnut color) to hold the channels in PDMS and the coverglass together.

**Figure 1–Figure supplement 2.**
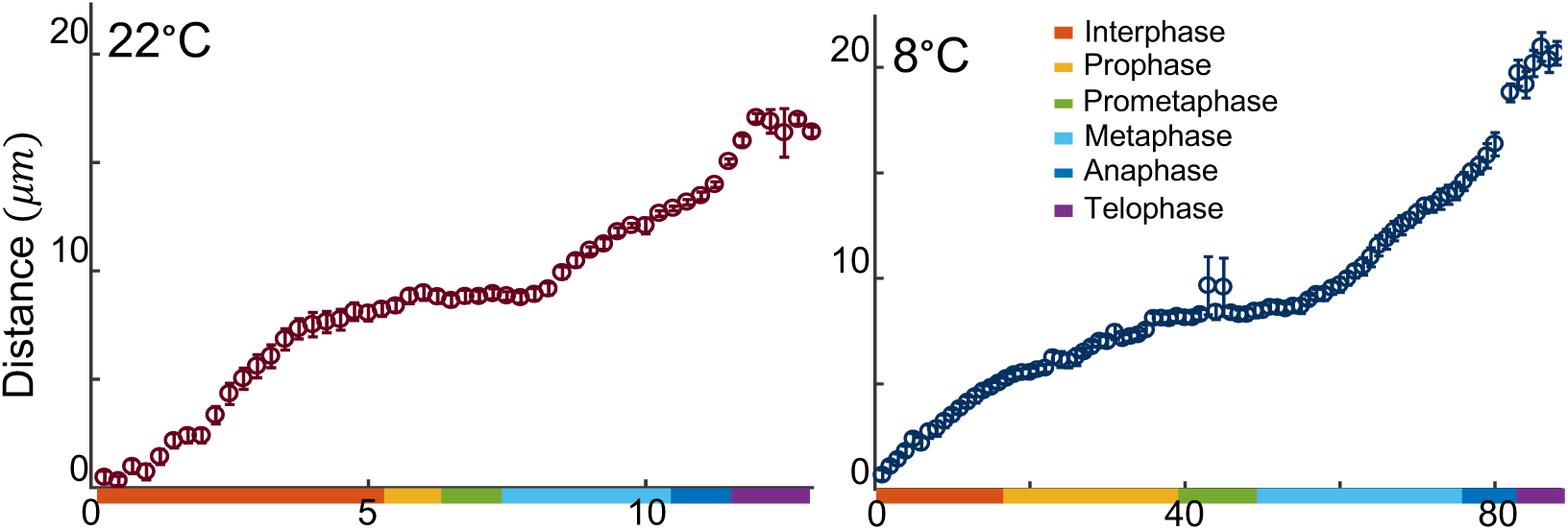
Average ± SEM for the distance between centrosomes was measured during NC11 for embryos developing at 22 and 8°*C*.

**Figure 1–Figure supplement 3.**
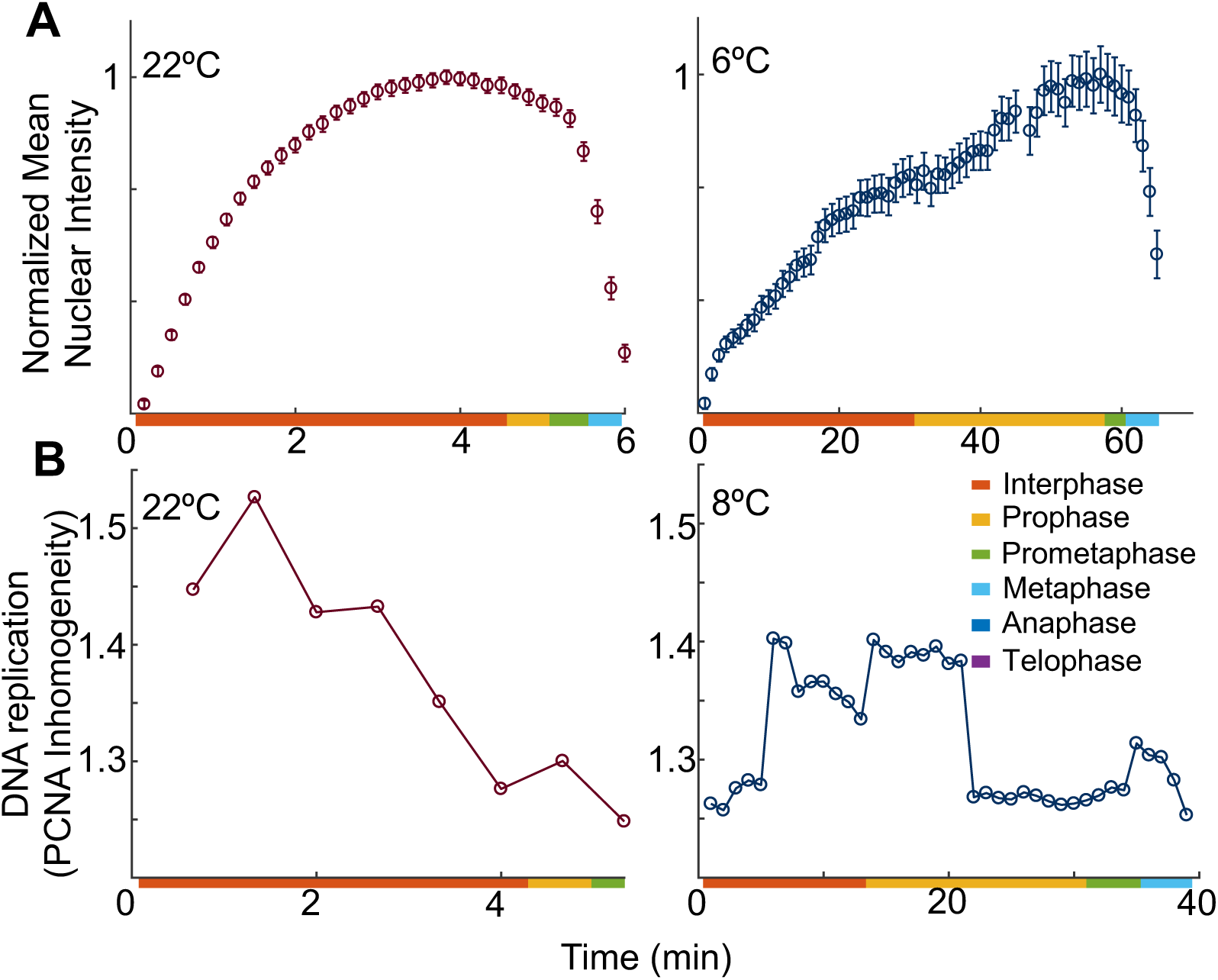
**A** Average ± SEM for nuclear intensity of a nuclear protein, Fibrillarin, at two different temperatures as a proxy for nuclear envelope breakdown. **B** A live reporter for DNA replication, PCNA, was used. During interphase, PCNA forms nuclear foci that are assumed to be DNA replication factories. The presence of these foci can be quantified as described in methods, as a proxy for DNA replication. All embryos are at NC11.

**Figure 1–Figure supplement 4.**
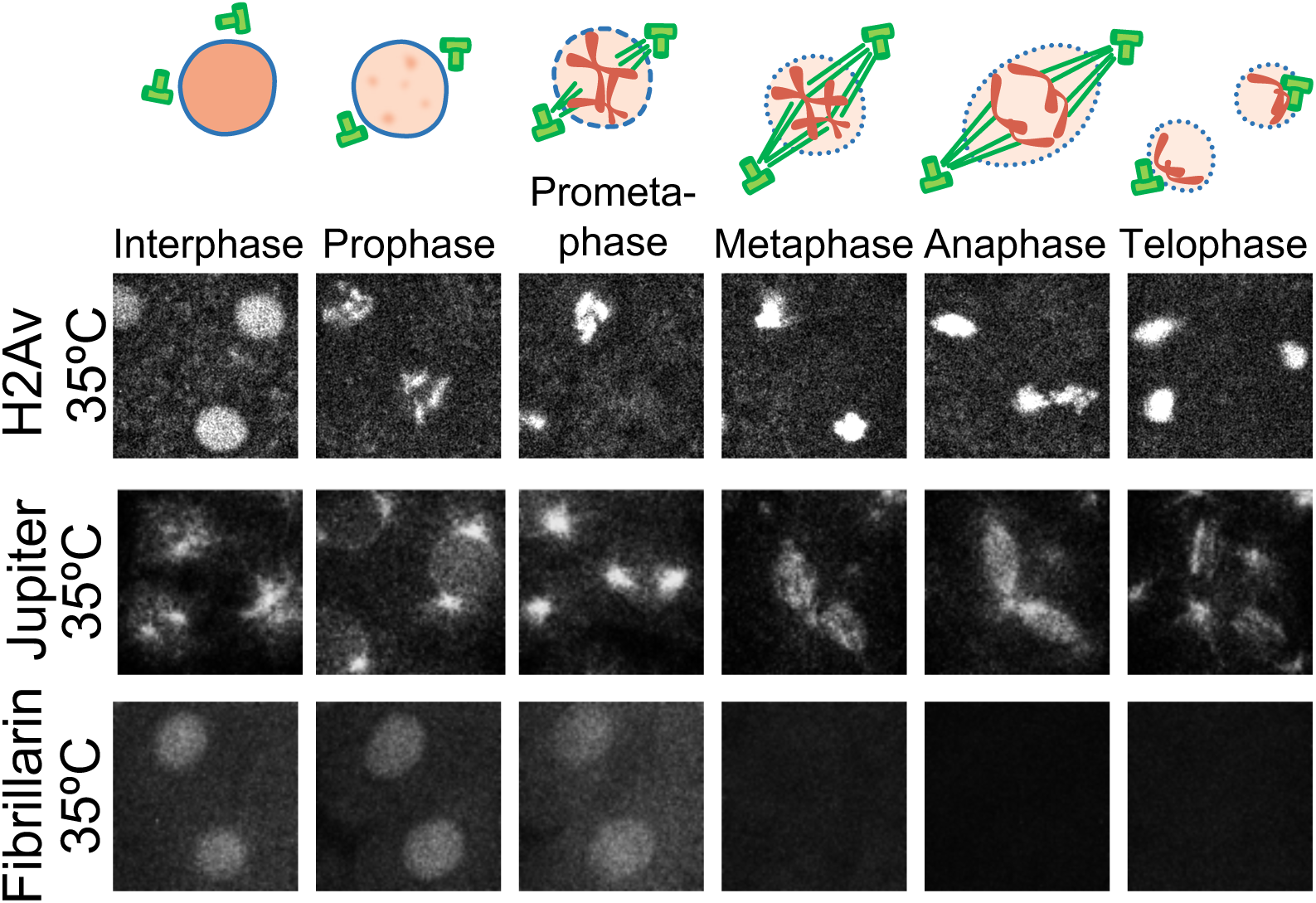
Fluorescently-tagged proteins are used to visualize different steps of NC11 for embryos developing at 35°C. Images are 25× 25*μm*.

**Figure 3–Figure supplement 1.**
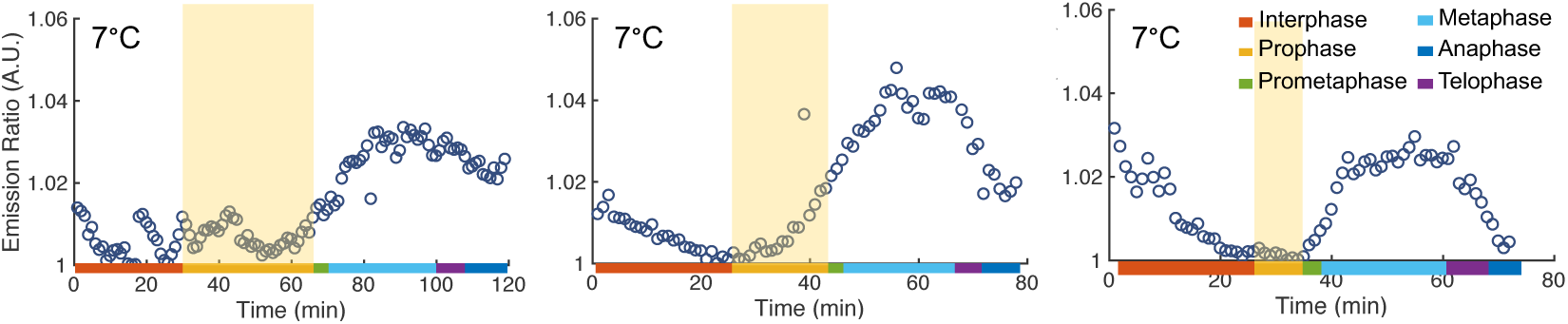
Emission ratio for the FRET sensor of Cdk1 activity of three embryos developing at 7°C. Emission ratio for embryos at room temperature was characterized extensively before (***Deneke et al., 2016***).

**Figure 3–Figure supplement 2.**
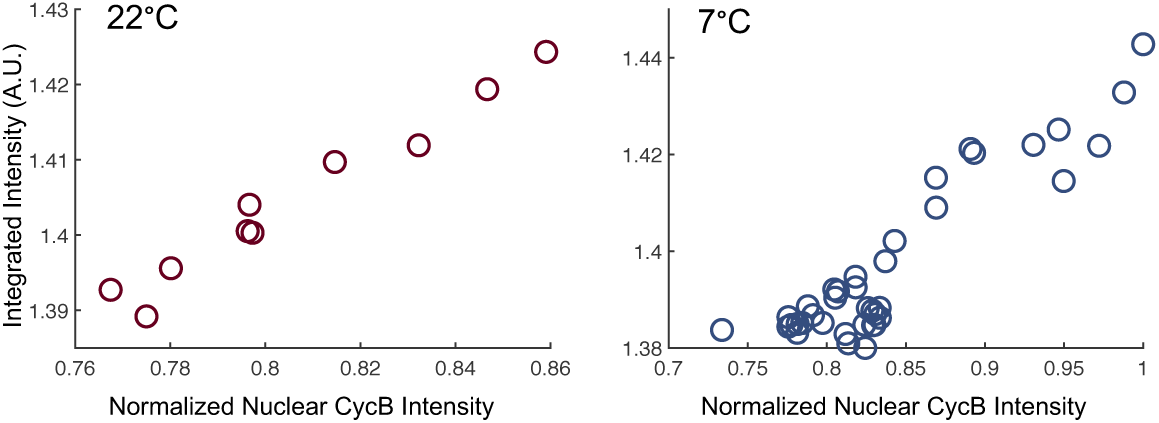
The rate of nuclear Cdk1 activity as detected by FRET sensor is plotted against the rate of nuclear accumulation of CycB. A linear relationship is expected if CycB accumulation is rate-limiting for the activity of Cdk1, and is observed at both temperatures. If the rate-limiting step for Cdk1 activity were after the nuclear accumulation of CycB, the expected outcome would have been for nuclear levels of CycB to reach equilibrium while Cdk1 activity would continue to increase. Therefore, our data supports a model in which CycB accumulation is rate limiting for Cdk1 activity. Data from embryos displayed in **Figure 3**A-B, and only includes the data-points were the reverse reactions (Cdk1 inactivation and CycB degradation) are negligible. Left: 22°C from 4-6.5 min, right: 7°C from 18-55 *min*.

## References

Adams RR, Maiato H, Earnshaw WC, Carmena M. Essential Roles of Drosophila Inner Centromere Protein (Incenp) and Aurora B in Histone H3 Phosphorylation, Metaphase Chromosome Alignment, Kinetochore Disjunction, and Chromosome Segregation. The Journal of Cell Biology. 2001 May; 153(4):865–880. http://dx.doi.org/10.1083/jcb.153.4.865, doi: 10.1083/jcb.153.4.865.

Arrhenius S. Quantitative laws in biological chemistry, vol. 1915. G. Bell; 1915.

Begasse ML, Leaver M, Vazquez F, Grill SW, Hyman AA. Temperature Dependence of Cell Division Timing Accounts for a Shift in the Thermal Limits of C. elegans and C. briggsae. Cell Reports. 2015; 10(5):647 – 653. http://www.sciencedirect.com/science/article/pii/S2211124715000078, doi: https://doi.org/10.1016/j.celrep.2015.01.006.

Blythe SA, Wieschaus EF. Establishment and maintenance of heritable chromatin structure during early *Drosophila* embryogenesis. eLife. 2016 nov; 5:e20148. https://doi.org/10.7554/eLife.20148, doi: 10.7554/eLife.20148.

Dell AI, Pawar S, Savage VM. Systematic variation in the temperature dependence of physiological and ecological traits. Proceedings of the National Academy of Sciences. 2011; 108(26):10591–10596. https://www.pnas.org/content/108/26/10591, doi: 10.1073/pnas.1015178108.

Denarier E, Fourest-Lieuvin A, Bosc C, Pirollet F, Chapel A, Margolis RL, Job D. Nonneuronal isoforms of STOP protein are responsible for microtubule cold stability in mammalian fibroblasts. Proceedings of the National Academy of Sciences. 1998; 95(11):6055–6060. https://www.pnas.org/content/95/11/6055, doi: 10.1073/pnas.95.11.6055.

Deneke VE, Melbinger A, Vergassola M, Talia SD. Waves of Cdk1 Activity in S Phase Synchronize the Cell Cycle in Drosophila Embryos. Developmental Cell. 2016; 38(4):399 – 412. http://www.sciencedirect.com/science/article/pii/S1534580716305160, doi: https://doi.org/10.1016/j.devcel.2016.07.023.

Dura JM. Stage dependent synthesis of heat shock induced proteins in early embryos of Drosophila melanogaster. Molecular and General Genetics MGG. 1981 Dec; 184(3):381–385. http://dx.doi.org/10.1007/BF00352509, doi: 10.1007/bf00352509.

Edgar BA, Sprenger F, Duronio RJ, Leopold P, O’Farrell PH. Distinct molecular mechanism regulate cell cycle timing at successive stages of Drosophila embryogenesis. Genes & Development. 1994; 8(4):440–452. http://genesdev.cshlp.org/content/8/4/440.abstract, doi: 10.1101/gad.8.4.440.

Falahati H, Pelham-Webb B, Blythe S, Wieschaus E. Nucleation by rRNA Dictates the Precision of Nucleolus Assembly. Current Biology. 2016; 26(3):277 – 285. http://www.sciencedirect.com/science/article/pii/ S0960982215015134, doi: https://doi.org/10.1016/j.cub.2015.11.065.

Falahati H, Wieschaus E. Independent active and thermodynamic processes govern the nucleolus assembly in vivo. Proceedings of the National Academy of Sciences. 2017; 114(6):1335–1340. https://www.pnas.org/content/114/6/1335, doi: 10.1073/pnas.1615395114.

Farrell JA, Shermoen AW, Yuan K, O’Farrell PH. Embryonic onset of late replication requires Cdc25 down-regulation. Genes & Development. 2012; 26(7):714–725. http://genesdev.cshlp.org/content/26/7/714.abstract, doi: 10.1101/gad.186429.111.

Ferree PL, Di Talia S. Chemical Waves in Embryonic Cell Cycles. Israel Journal of Chemistry. 2018; 58(6-7):714–721. https://onlinelibrary.wiley.com/doi/abs/10.1002/ijch.201700144, doi: 10.1002/ijch.201700144.

Gavet O, Pines J. Progressive Activation of CyclinB1-Cdk1 Coordinates Entry to Mitosis. Developmental Cell. 2010; 18(4):533 – 543. http://www.sciencedirect.com/science/article/pii/S1534580710001115, doi: https://doi.org/10.1016/j.devcel.2010.02.013.

Giet R, Glover DM. Drosophila Aurora B Kinase Is Required for Histone H3 Phosphorylation and Condensin Recruitment during Chromosome Condensation and to Organize the Central Spindle during Cytokinesis. The Journal of Cell Biology. 2001 Feb; 152(4):669–682. http://dx.doi.org/10.1083/jcb.152.4.669, doi: 10.1083/jcb.152.4.669.

Gillooly JF, Brown JH, West GB, Savage VM, Charnov EL. Effects of Size and Temperature on Metabolic Rate. Science. 2001; 293(5538):2248–2251. https://science.sciencemag.org/content/293/5538/2248, doi: 10.1126/science.1061967.

Heim A, Rymarczyk B, Mayer TU. Regulation of Cell Division. Vertebrate Development. 2016 Dec; p. 83–116. http://dx.doi.org/10.1007/978-3-319-46095-6_3, doi: 10.1007/978-3-319-46095-6_3.

Karpova N, Bobinnec Y, Fouix S, Huitorel P, Debec A. Jupiter, a new Drosophila protein associated with microtubules. Cell Motility. 2006; 63(5):301–312. https://onlinelibrary.wiley.com/doi/abs/10.1002/cm.20124, doi: 10.1002/cm.20124.

Kuntz SG, Eisen MB. Drosophila Embryogenesis Scales Uniformly across Temperature in Developmentally Diverse Species. PLOS Genetics. 2014 04; 10(4):1–15. https://doi.org/10.1371/journal.pgen.1004293, doi: 10.1371/journal.pgen.1004293.

Lindqvist A, Rodríguez-Bravo V, Medema RH. The decision to enter mitosis: feedback and redundancy in the mitotic entry network. Journal of Cell Biology. 2009 04; 185(2):193–202. https://doi.org/10.1083/jcb.200812045, doi: 10.1083/jcb.200812045.

Lucchetta EM, Lee JH, Fu LA, Patel NH, Ismagilov RF. Dynamics of Drosophila embryonic patterning network perturbed in space and time using microfluidics. Nature. 2005 Apr; 434(7037):1134–1138. http://dx.doi.org/10.1038/nature03509, doi: 10.1038/nature03509.

McCleland ML, O’Farrell PH. RNAi of Mitotic Cyclins in Drosophila Uncouples the Nuclear and Centrosome Cycle. Current Biology. 2008; 18(4):245 – 254. http://www.sciencedirect.com/science/article/pii/S0960982208000882, doi: https://doi.org/10.1016/j.cub.2008.01.041.

Morgan DO. The cell cycle: principles of control. New Science Press; 2007.

Novak ZA, Wainman A, Gartenmann L, Raff JW. Cdk1 Phosphorylates Drosophila Sas-4 to Recruit Polo to Daughter Centrioles and Convert Them to Centrosomes. Developmental Cell. 2016; 37(6):545 – 557. http://www.sciencedirect.com/science/article/pii/S1534580716303355, doi: https://doi.org/10.1016/j.devcel.2016.05.022.

Pomerening JR, Sontag ED, Ferrell JE. Building a cell cycle oscillator: hysteresis and bistability in the activation of Cdc2. Nature Cell Biology. 2003 Mar; 5(4):346–351. http://dx.doi.org/10.1038/ncb954, doi: 10.1038/ncb954.

Rabinowitz M. Studies on the cytology and early embryology of the egg of Drosophila melanogaster. Journal of Morphology. 1941; 69(1):1–49. https://onlinelibrary.wiley.com/doi/abs/10.1002/jmor.1050690102, doi: 10.1002/jmor.1050690102.

Raven JA, Geider RJ. Temperature and algal growth. New Phytologist. 1988; 110(4):441–461. https://nph.onlinelibrary.wiley.com/doi/abs/10.1111/j.1469-8137.1988.tb00282.x, doi: 10.1111/j.1469-8137.1988.tb00282.x.

